# Refutation of traumatic insemination in the *Drosophila bipectinata* species complex: Hypothesis fails critical tests

**DOI:** 10.1101/2021.08.16.456442

**Authors:** Michal Polak, Shane F. McEvey

**Affiliations:** Department of Biological Sciences, University of Cincinnati, Cincinnati, OH 45221-0006 United States of America; Australian Museum Research Institute, Australian Museum, 1 William Street, Sydney NSW 2010, Australia

**Keywords:** *Drosophila bipectinata* species complex, traumatic insemination hypothesis, copulation, aedeagal lateral processes, genital claws, anchoring, sperm delivery

## Abstract

Traumatic insemination (TI) is a rare reproductive behaviour characterized by the transfer of sperm to the female via puncture wounds inflicted across her body wall. Here, we challenge the claim made by Kamimura (2007) that males of species of the *Drosophila bipectinata* complex utilize a pair of claw-like processes (“claws”) to traumatically inseminate females: the claws are purported to puncture the female body wall and genital tract, and to inject sperm through the wounds into the genital tract, bypassing the vaginal opening, the route of sperm transfer occurring in other *Drosophila*. This supposed case of TI is widely cited and featured in prominent subject reviews. We examined high-resolution scanning electron micrographs of the claws and failed to discover any obvious “groove” for sperm transport. We demonstrated that sperm occurred in the female reproductive tract as a single integrated unit when mating flies were experimentally separated, inconsistent with the claim that sperm are injected via paired processes. The aedeagus in the *bipectinata* complex was imaged, and shown to deliver sperm through the vaginal opening. Laser ablation of the sharp terminal ends of the claws failed to inhibit insemination. The results refute the claim of TI in the *Drosophila bipectinata* species complex.

## 1. Introduction

Traumatic insemination (TI) is a form of mating behaviour during which males employ specialized “devices”, such as spines and stylets, to puncture the female body wall and transfer sperm through the wound(s) (Lange et al., 2013). This extraordinary behaviour is distinguished from other forms of “traumatic mating”, where only non-sperm components of the ejaculate, or no ejaculate at all, transfer to the female through male-inflicted wounds (Blanckenhorn et al., 2002; Siva-Jothy, 2009; Lange et al., 2013; Reinhardt et al., 2015).

Though rare, TI *sensu stricto* has arisen independently in a number of animal groups (Reinhardt et al., 2015), and the evolutionary drivers of this unusual form of insemination are of considerable interest and the focus of ongoing debate (Eberhard 1985, 1996; Arnqvist and Rowe, 2005; Tatarnic and Cassis, 2010; Lange et al., 2013; Tatarnic et al., 2014; Dougherty et al., 2017; Brand et al., 2021). The identification of *bona fide* cases of TI in animals will be important not only for accurately documenting the taxonomic distribution of this remarkable form of mating (Reinhardt et al., 2015), but also for facilitating the comparative method to test hypotheses about the selective pressures favoring its emergence in evolution (Futuyma and Kirkpatrick, 2017, p. 69). Distinguishing the various forms of traumatic mating also lies at the heart of our ability to predict specific selective challenges females may face during mating, and hence to interpret immunological, anatomical, and behavioural features as potential counter-adaptations to variable forms of male-induced harm in the broader context of sexual conflict theory (Johnstone and Keller, 2000; Hosken et al., 2003; Morrow et al., 2003; Arnqvist and Rowe, 2005; Rönn et al., 2007; Siva-Jothy, 2009; Dougherty et al., 2017).

Among terrestrial arthropods, TI is particularly prevalent in the hemipteran infraorder Cimicomorpha, which includes the well-studied human bed bug, *Cimex lectularius*. Males of this species pierce the ventral surface of a female’s abdomen with a curved, needle-like, hollow stylet (or paramere) (Usinger, 1966) that possesses a sizeable pore near its tip (Fig. 1), through which sperm and seminal fluid are injected into the female (Davis, 1956; Carayon, 1966). Cases of TI have also been convincingly demonstrated in the plant bug genus *Coridromius* (Tatarnic & Cassis, 2010), the spider *Harpactea sadistica* (Řezáč, 2009), and the marine flatworm *Pseudocerus bifurcus* (Michiels and Newman, 1998; and see Brand et al., 2021).

**Figure 1.**
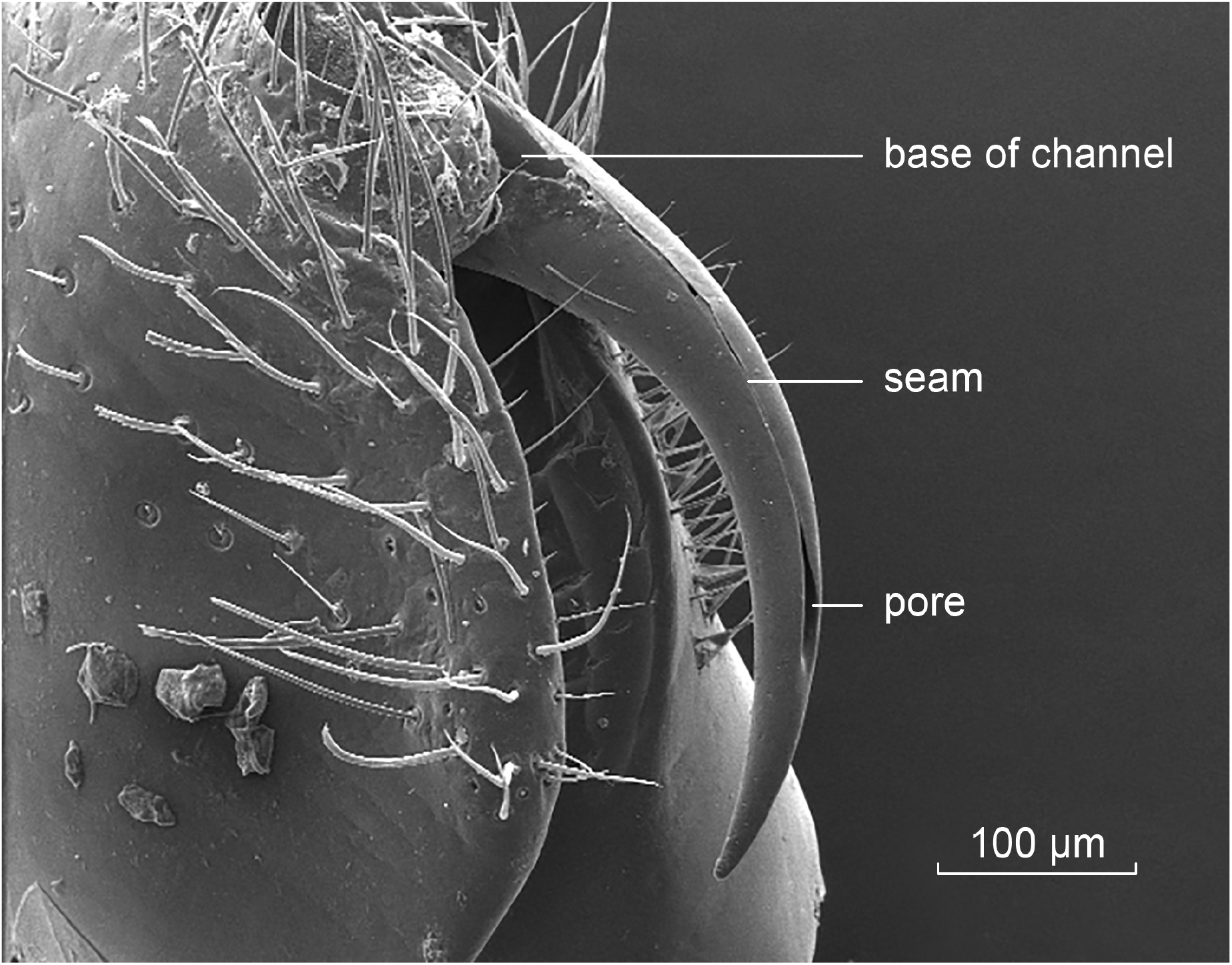
Scanning electron micrograph (200×) of the curved and tapered copulatory organ (the paramere, Usinger, 1966) of the bed bug, *Cimex lectularius* L. During mating, the paramere can be seen in real time to puncture the female abdominal integument (Usinger, 1966). The channel (groove) of the paramere extends from its base to the subterminal pore (c. 65 × 10 µm); the aedeagus is everted through the channel and ejaculate injected into the female (Davis, 1956; Usinger, 1966).

It has been claimed by Kamimura (2007) (henceforth K2007) that TI occurs in species of the *Drosophila bipectinata* complex, a small taxonomic grouping of four very similar species in the *ananassae* subgroup of the *melanogaster* species group, that includes: *D. bipectinata* Duda, 1923; *D. parabipectinata* Bock, 1971; *D. malerkotliana* Parshad and Paika, 1965; and *D. pseudoananassae* Bock, 1971 (Bock, 1971). Specifically, males of these species are purported to use a pair of claw-like phallic structures, called “basal processes” by K2007, but which we call aedeagal lateral processes following Rice et al. (2021), as explained below, to pierce the female body wall and reproductive tract, and inject sperm into her reproductive tract through the wound sites. If true, then TI in this complex would be an astonishing evolutionary innovation within the genus, family and order.

A wide range of species of *Drosophila* and related genera have been used to investigate dipteran reproductive biology in detail. This body of knowledge both underscores the astounding nature of the TI claim and provides a thorough understanding of the “typical” route of sperm transfer, storage and use, which is particularly well understood for the model organism, *D. melanogaster*. In this species, mating lasts approximately 20 min during which the male and female attach through the union and integration of various male and female genital structures (Eberhard and Ramirez, 2004; Jagadeeshan and Singh, 2006; Mattei et al., 2015). The intromittent organ in males is the aedeagus, and the ejaculate, comprised of sperm and seminal plasma, pass via the tip of the aedeagus into the female uterus (bursa) via her gonopore (Bairati, 1968; Fowler, 1973; Manier et al., 2010; Mattei et al., 2015). The transferred ejaculate fills and can considerably swell the bursa in this and other species (Patterson, 1946; Lefevre and Jonsson, 1962; Markow and Ankney, 1988; Alonso-Pimentel et al., 1994). Upon dissection of the female after the termination of copulation, the ejaculate can readily be visualized intact by teasing the bursa open and releasing the sperm mass associated with its “waxy” plug material (Alonso-Pimentel et al., 1994; Pitnick and Markow, 1994; Polak et al., 1998). As sperm entry into storage within the female requires some time to begin (approximately 4 minutes in *D. melanogaster*) after the end of copulation (Manier et al., 2010), the dissected sperm mass may be discerned intact with its full complement of sperm, so long as it is dissected from the bursa quickly after copulation (Pitnick and Markow, 1994; Manier et al., 2010; Polak and Rashed, 2010; Tyler et al., 2021).

In K2007’s study, adult males were allowed to ingest food containing rhodamine-B fluorescent dye, and were mated to virgin females. Mating pairs were flash frozen in liquid nitrogen 5 minutes after the onset of copulation (uninterrupted copulations last, on average, approximately 10.6 min in *D. bipectinata*, Polak et al., 2021, and see below), and abdomens of coupled pairs removed, mounted and observed. A laser scan micrograph showing two areas of coloration adjacent to the tips of the basal processes (figure 1 (c) (ii) in K2007) was evidence put forward for the occurrence of TI. On the basis of these images, Kamimura claimed that the two claw-like processes pierce the female’s body wall, and transfer sperm through the wounds into the female reproductive tract, bypassing the opening of the vagina (gonopore). The following is the relevant excerpt from K2007 (pp. 403–404): “TI clearly occurs in the *bipectinata* complex, as the basal processes pierce the pockets during copulation and sperm is ejaculated through the wounds but not through the genital orifice…”. No direct evidence for the passage of sperm via such a route was provided. The following statement likewise gave us pause (p. 404): “The basal processes of this group have a groove on the dorsal surface which may transport semen.” The problem also here is that no visual evidence for such a groove was presented.

Another source of our skepticism regarding K2007’s claim comes from our own observations of the ejaculated sperm mass in *D. bipectinata* made previously (Polak and Rashed, 2010; Tyler et al., 2021). Upon dissection from the female bursa immediately after copulation, the sperm in *D. bipectinata*, which are 1.63 mm long (Tyler et al., 2021) and, for reference, slightly shorter (by c. 12%) than in *D. melanogaster* (1.85 mm, (Manier et al., 2013)), invariably occur as a *single and strongly integrated mass* in association with characteristic “waxy” plug material, similar in appearance to that seen in *D. melanogaster*. This observation challenges the TI hypothesis of insemination via paired “basal processes”, because if TI occurs we would expect the injected sperm mass within the bursa immediately after copulation to be discernable as two more-or-less distinct units, but we have consistently observed only a single, highly integrated mass (Tyler et al., 2021). The sperm tails in *D. bipectinata* are long and the sperm mass consequently is difficult to disentangle (Tyler et al., 2021), so the idea that two separate masses injected via the paired “basal processes” would then dynamically coalesce during mating, or shortly thereafter, and form a single homogeneous mass with associated plug material, seems unlikely.

Here, we challenge the claim that TI occurs in the *bipectinata* complex. Our results provide observational and direct experimental evidence contradicting such a claim: the evidence supports sperm transfer occurring via the route of sperm delivery in other *Drosophila*, that is, through the female gonopore into the reproductive tract. In the present section we first offer a reinterpretation of the male phallic architecture of species within the *bipectinata* complex, following the standardized nomenclature proposed for *D. melanogaster* (Rice et al., 2019) and refined for a wider group of species following developmental studies (Rice et al., 2021). On the one hand, we agree with K2007 that the paired claw-like structures, the purported piercing organs of TI, are not a “bifid aedeagus”, a terminology adopted by earlier authors. Okada ((Okada, 1954), pl. 3, fig. 14) illustrated the *D. bipectinata* male terminalia and labelled the paired structures “aedeagus”, and Parshad and Paika used the term “bifid” in describing the same paired structures (Parshad and Paika, 1964). Bock (1971) used the descriptor “aedeagus bifid and bare” to distinguish the *bipectinata* complex, and Bock and Wheeler (1972) and Gupta (1973) followed this usage. On the other hand, we also agree with Rice et al. ‘s (2021) suggestion that the claw-like structures should be termed “aedeagal lateral processes”, rather than K2007’s “basal processes”. Rice et al. (2021) showed that the claws have a distinct developmental origin, deriving from the *lateral* portions of the central primordium of the phallus observable during metamorphosis in the pupa. Since most of the cells of the central primordium normally give rise, in related *Drosophila* subgroups, to the aedeagus (not to postgonites, pregonites, or postgonal sheath) the term *aedeagal lateral process* was proposed by Rice et al. (2021), a term we adopt here.

As a first step in our test of the TI hypothesis, we examined scanning electron micrographs of the aedeagal lateral processes (claws) at varying orientations and magnifications to search for the so-called “groove”—the alleged conduit for sperm delivery. We then addressed two additional predictions. The first is that the sperm mass should be observable as two more-or-less distinct units upon transfer to the female, as mentioned above. To this end, we dissected and examined the ejaculated sperm masses extracted from the female reproductive tract both immediately after the terminus of full-length, uninterrupted copulations, and after pairs were interrupted 6–8 min after the onset of coupling. Reproductive structures of both sexes were also examined after copulation interruption to elucidate the path of sperm transfer to the female. Finally, we used ultraprecise laser surgery (Polak and Rashed, 2010) to ablate the terminal ends of both claws in individual males, thus eliminating their pointed tips. If the claws serve to transfer sperm by piercing across the female’s body wall and reproductive tract, then males with surgically ablated piercing devices should fail to transfer sperm.

## 2. Material and methods

### (a) Source and culture of flies

*Drosophila bipectinata* Duda and *D. parabipectinata* Bock cultures were established with field-caught flies captured from the surface of fallen fruits in Taiwan (25°2′30.24″ N, 121°36′39.37″ E). A *D. malerkotliana* Parshad and Paika culture was established with flies from Thailand (8°54’22.24”N 98°31’43.51”E). Flies were cultured in half-pint glass bottles on standard cornmeal-agar medium within an environmental chamber under controlled light and temperature conditions (12h light (24°C):12h dark (22°C)). Adult virgin flies for mating trials were harvested under a light stream of humidified CO_2_ within 8 h of emergence, and housed in groups of 10–15 flies in 8-dram food vials containing cornmeal-agar medium. All flies used in mating trials were 4–6 d old. Material for morphological examination and imaging of genitalia were also derived from ethanol-preserved specimens held in the Australian Museum, Sydney (AMS K.380306–07).

### (b) Laser surgery

The laser surgical protocol is described in detail elsewhere (Polak and Rashed, 2010). Briefly, young males (< 24 h of age) were anesthetized with CO_2_ in an acrylic chamber with a thin glass bottom. The male was positioned ventral side down in the chamber, so the external genitalia were visible from below and accessible to the laser light. The chamber was mounted on a Prior (Rockland, MA, USA) H117 motorized stage fitted to an Olympus (Center Valley, PA, USA) IX71 inverted light microscope. Individual pulses of light (λ=532 nm) from a Vector 532-1000-20 Q-switched laser (Coherent, Santa Clara, CA, USA) focused through an Olympus UPlanApo 20x objective were used to ablate 1/4 to 1/3 of both lateral processes (claws) of individual males. After surgery, the fly was gently aspirated out of the chamber, and allowed to recover in groups of 3–5 males in food vials for at least 3 d until individually paired with virgin females. Uncut control males were treated identically to that as above, except that 1–2 large bristles near the apex of the abdomen were laser-ablated on both sides of the body; their claws were untouched by laser light. These constitute the co-called “surgical control” group used in previous work (Grieshop and Polak, 2012, 2014; Rodriguez**-**Exposito et al., 2020).

### (c) Mating trials and dissections

All mating trials were conducted in the morning between 8:00 am (lights on) and 11:00 am. Virgin males (4–6 d of age) were each individually paired with a virgin female (3–4 d old). The onset and termination of copulation were recorded, and copulation duration was taken as the difference between these time points. Immediately after the termination of copulation (when the male dismounted), the female was killed with ether fumes and dissected in a drop of phosphate-buffered physiological saline (PBS) under an Olympus SZX12 stereomicroscope (Olympus Corp. of the Americas, Center Valley, PA, USA). The female bursa was gently teased open using fine biology-grade forceps. As the sperm mass was released into the saline, it was ascertained whether it was in the form of a single mass or > 1 mass. On two separate mornings, copulations were interrupted using a fine paintbrush to separate the pair. Immediately after the pair was separated, the female was dissected and the sperm mass, if present, was released from the bursa and examined, as above. A total of 12 copulations with different individuals were interrupted between 6 and 8 minutes after the start of mating. One copulation was interrupted at 4 min; no ejaculate could be detected within the female.

### (d) Imaging phallic structures

Genitalia were dissected from alcohol-preserved specimens, or from fresh material from culture, using a common procedure. Under a stereomicroscope, reproductive structures from fresh material for examination and imaging were dissected or extruded in a few drops of 1× phosphate buffered saline (PBS) on a depression slide and imaged immediately. For male genitalic structures, genitalia were dissected into 70% ethanol and teased free of attached pieces of exoskeleton and soft tissue using fine tweezers and dissecting probes. The specimen then was gently boiled in 1N KOH for ≈ 8 min to dissolve soft tissue and to improve observation of the hard parts. Digital images of fresh material and boiled genital structures were captured with a Leica M205 Stereomicroscope (Leica Microsystems, Buffalo Grove, IL, USA). The light microscope images were used for 1) imaging freshly dissected reproductive structures (sperm masses, female uterus, male aedeagus and female oviscape), 2) visualizing, describing and annotating phallic and periphallic structures, and 3) for confirming the integrity of laser-surgical manipulations. For scanning electron micrograph (SEM) acquisition, specimens of fly genitalia were dissected and treated with KOH as above. They were then rinsed in distilled water, air-dried, mounted on conductive carbon adhesive tabs atop an aluminum post, adjusted for proper orientation, sputter coated with gold-palladium film, and imaged with a SCIOS Dual-Beam Scanning Electron Microscope (ThermoFisher, Waltham, MA, USA). SEMs were used primarily for examining the claws for possible sperm conduit architecture, and for producing exemplars of the laser cuts. To check for sperm conduits, multiple images of the same structure were taken at different magnifications (typically between 350–3500×) and different orientations effected by motorized tilting of the specimen within the microscope chamber. Genitalic preparations of a total of 28, 29, and 3 different individuals of *D. parabipectinata, D. bipectinata* and *D. malerkotliana*, respectively, were imaged, and a total of 166 SEMs were examined and archived.

## 3. Results

### (a) Phallic architecture of the *bipectinata* species complex

The phallic structures of the *bipectinata* complex are represented here by *D. bipectinata* and *D. parabipectinata*; the same, or very similar, morphology is present in the other two species of the *bipectinata* complex (Bock, 1971). Naming the parts of the copulatory apparatus is difficult because they are exceptionally evolutionary labile, their homologies obscure (but see Rice et al., 2021), and their function often speculative.

The male hypandrium of the *bipectinata* complex species was imaged within the body (Fig. 2A). The aedeagal lateral processes (claws) (Fig. 2) are articulated with the apex of the phallapodeme (articulation arrowed in Fig. 2D). The aedeagal lateral processes are large, curved, apically pointed and bare (Fig. 2C, D; Fig. 3A–E), approximately 90 µm in length from base to tip. They are bilaterally symmetrical, and arise from the lateral portions (“shoulders”) (Fig. 2A, D, E), not the center, of the apex of the phallapodeme (this comports with the ontogeny described by Rice et al., 2021).

**Figure 2.**
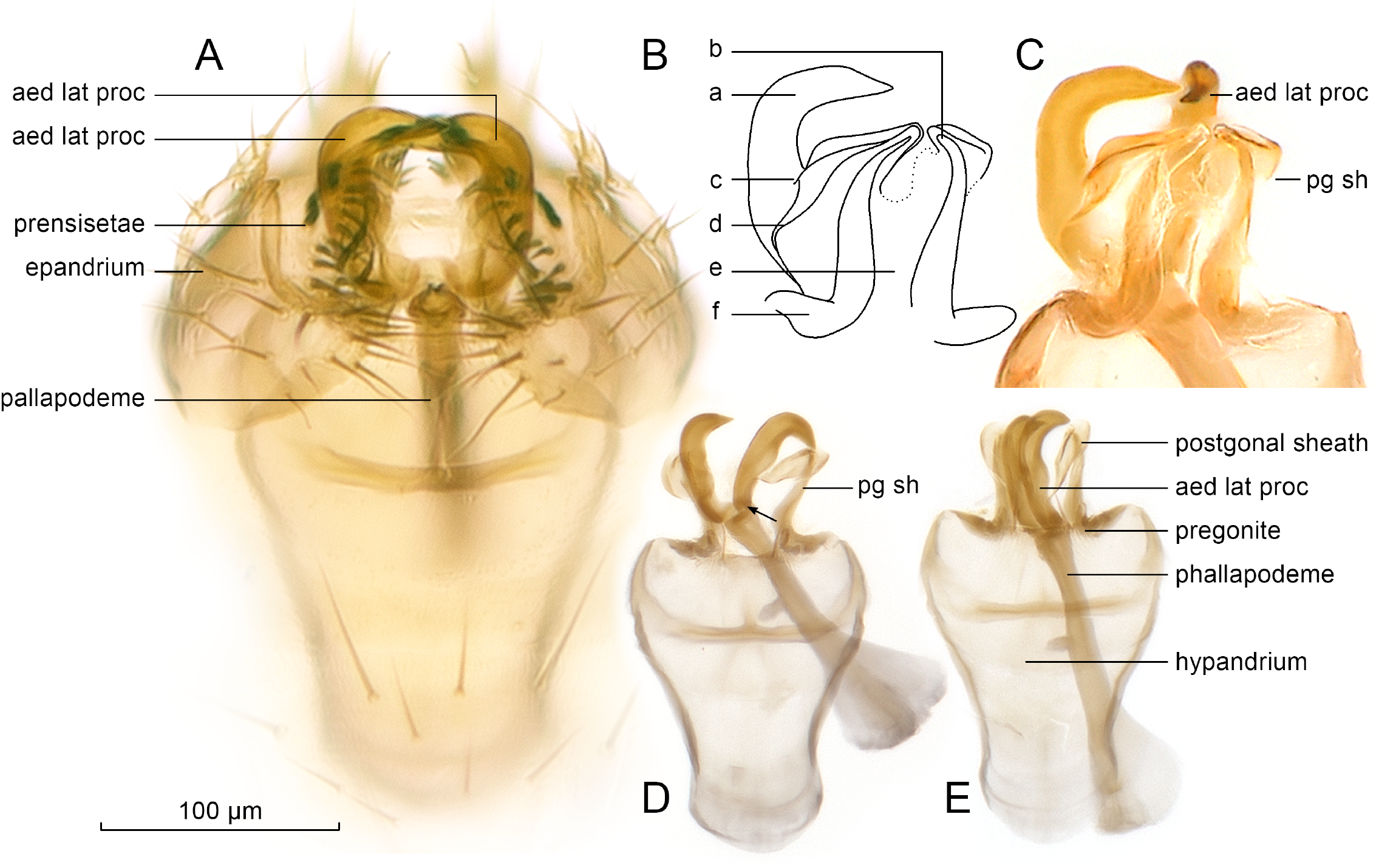
Light microscope images of the phallic structures of the *bipectinata* complex. **(A)** Posterior end of the undissected male body viewed from below; the hypandrium is inside the body, surrounded by the epandrium. **(B, C)** Drawing and magnified view, after dissection, of the posterior end of the hypandrium, showing the aedeagal lateral processes (aed lat proc) and the postgonal sheath (pg sh). The sheath surrounds the phallic structures dorsally; it is largely membranous (e) but ridges and thickened processes (b, c, d, f) are evident within. **(D, E)** Views of the hypandrium in its entirety. Images show the aedeagal lateral processes and postgonal sheath in two phallapodeme orientations (projected and withdrawn, respectively), as well as the articulation of the aedeagal lateral process (arrowed in D) at the posterior end of the phallapodeme. Specimens: (A) *D. bipectinata* (Cape Tribulation, Australia | 16.104°S 145.455°E | 2011 | M. Polak & S.F. McEvey); (B, C) *D. bipectinata* (Taipei, Taiwan | 25°2’30.24”N, 121°36’39.37”E | 2017 | M. Polak); (D, E) *D. parabipectinata* (Christmas Island [nr Java] | 10°30’S 105°35’E | 2003 | S.F. McEvey et al. [Australian Museum]).

**Figure 3.**
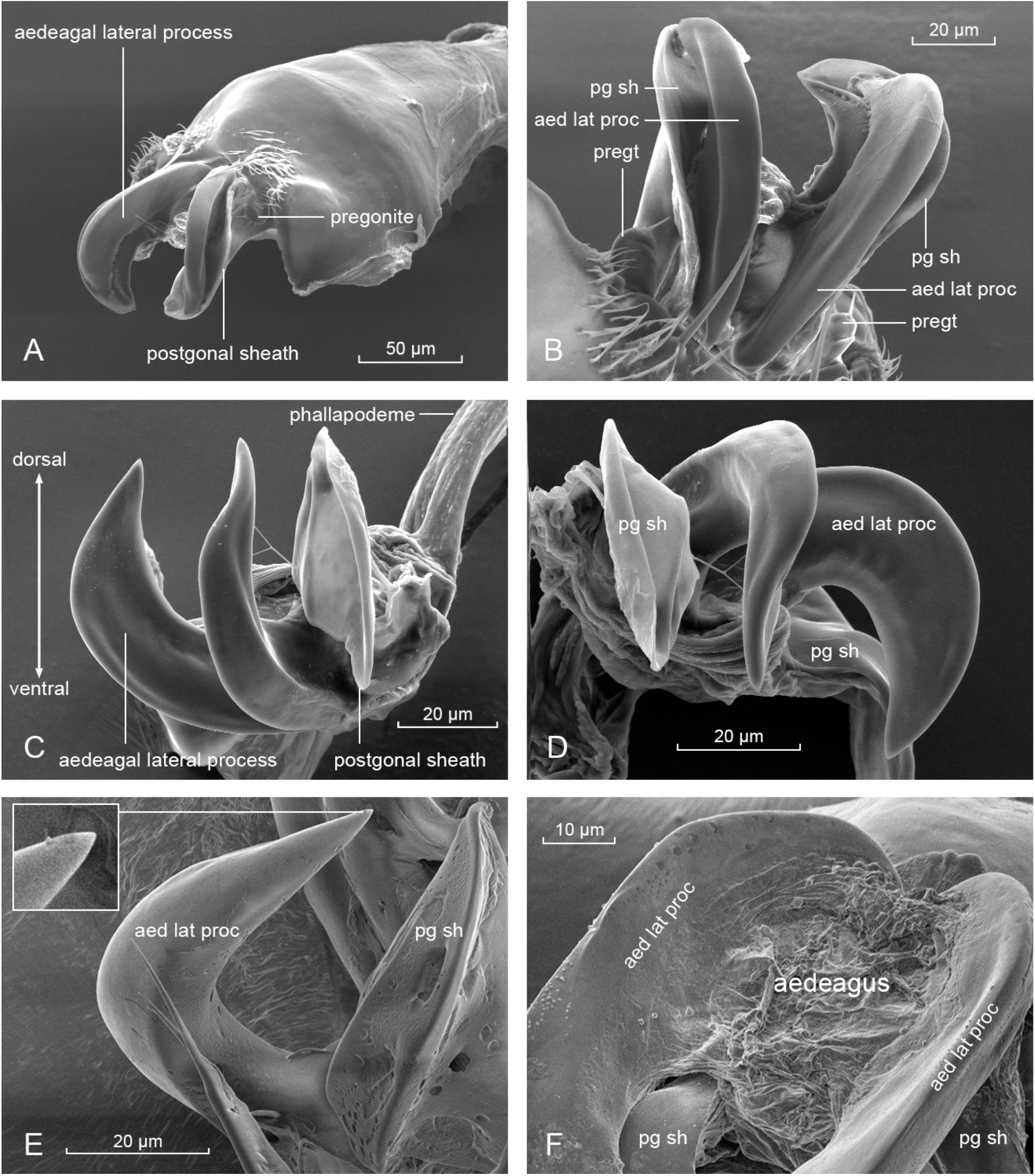
Scanning electron micrographs of the aedeagal lateral processes (aed lat proc) in the *bipectinata* complex. Aedeagal lateral processes were imaged at different orientations and magnifications to allow detailed examination of all surfaces, which failed to reveal purported conduits (grooves or channels) for sperm transport. **(A)** Ventral surface of hypandrium with the pair of aedeagal lateral processes, sheathed in the postgonal sheath (pg sh); the small pregonite (pregt) with apical setae arises from the gonocoxite. **(B)** Strong, rounded, bare, ventral surfaces of the aedeagal lateral processes with the postgonal sheath extending beyond (and possibly protecting) the sharp tips. Pregonite (pregt) with apical sensilla. **(C)** Smooth, bare, seamless, ventral and lateral surfaces of the aedeagal lateral processes; no gonopore (phallotrema) present; thickened edges and internal “ribs” of the postgonal sheath visible (see also D, and Fig. 2B, C). **(D)** Ventral and lateral faces of the aedeagal lateral processes, in relation to the lobes of the postgonal sheath. **(E)** Surfaces of aedeagal lateral process. Inset shows intact tip, which occasionally carries a lesion, possibly an abrasion (not shown). **(F)** Reticulated mat of tissue between the bases of the aedeagal lateral processes, interpreted to be the collapsed aedeagus (see Fig. 5A, B). Specimens: (A–D) *D. parabipectinata*, (E, F) *D. bipectinata* (Taipei, Taiwan | 25°2’30.24”N, 121°36’39.37”E | 2017 | M. Polak).

We see no connection between the base of the pregonites and the claws (Fig. 2D, E), which confirms that the claws are also not “basal extensions” of the pregonites that exist in closely allied species in the *D. ananassae* complex (Bock and Wheeler, 1972; McEvey and Schiffer, 2015); in such species a very clear nexus exists between the pregonite and a large structure that curves and extends caudally from its base, the basal extension (McEvey and Schiffer, 2015).

The pregonites are small, rounded and J-shaped (Fig. 2E), with very small apical setae or “pregonal bristles” (Rice et al., 2019; Fig. 3A, B). They are derived from ventral primordial cells and are, therefore, developmentally separate from the aedeagal lateral processes and the postogonal sheath. The postgonal sheath (*sensu* Rice et al., 2021) is membranous, folds and bends freely, and it is loosely symmetrical, lobe-like and dorsal to the aedeagal lateral processes (Fig. 3A–D). When viewed via light microscopy it is largely membranous and transparent (Fig. 2C), with hardened outer ridges and leaf-like structure connected to the base of each claw (Fig. 2B–E). The postgonal sheath arises from the dorsolateral primordial cells, which, in other species, develop into postgonites (posterior parameres) (Rice et al., 2021). Postgonites are absent in the four species of the *bipectinata* complex.

### (b) The traumatic insemination hypothesis

We examined SEMs of the claws, including their dorsal, ventral and lateral surfaces (Fig. 3). The claws are bare and smooth on all surfaces, and when the phallapodeme is not extended, they are “cloaked” by the sheath dorsally (Fig. 3A, B). Critically, we could identify no channel, groove or fold, medially, laterally, dorsally or ventrally, on the claws that would function as a conduit for sperm. In some preparations (not shown) we observed the tip of one or both claws to have an irregular depression or lesion (typically ≤ 1 µm in diameter), which could be a result of abrasion given the often irregular (torn) edges of these spots.

When females were dissected immediately upon the termination of copulation, the sperm within the bursa invariably occurred as a single mass (Fig. 4A). Out of a total of 12 matings interrupted at 6–8 min after the start of mating, 11 produced a sperm mass within the female; in all these cases the sperm likewise occurred as a single mass within the bursa (Fig. 4B). In 3 cases, the sperm mass was small and appeared irregular in shape, amorphous, not smoothly oval or rounded, but nevertheless unquestionably as a single unit.

**Figure 4.**
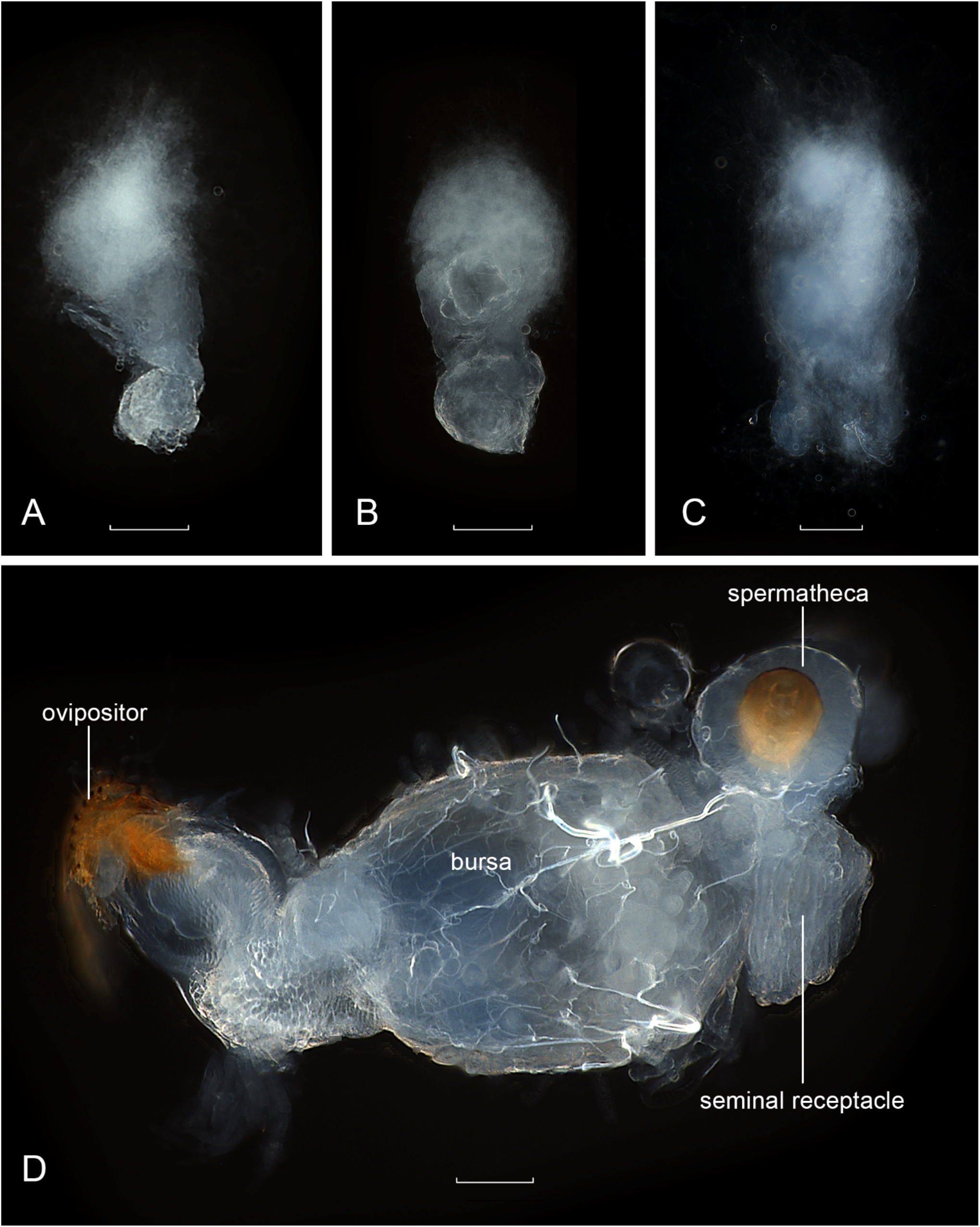
Light microscope images exemplifying the single sperm mass in *D. bipectinata*. Masses were dissected intact from the female reproductive tract **(A)** immediately after the end of a full-length, uninterrupted copulation, and **(B)** immediately after the pair was experimentally separated at 8 min or less after the onset of genital coupling; uninterrupted copulation duration in this species is on average about 10.6 min (Polak et al., 2021). At the base of each mass, the gelatinous (“waxy”) component of the ejaculate is clearly visible, which is also present in *D. melanogaster* and in other species to varying degrees of expression (Bairati and Perotti, 1970; Alonso-Pimentel et al., 1994; Polak et al., 1998; Manier et al., 2010). **(C)** Sperm mass dissected from the bursa after copulation with a male with both aedeagal lateral process tips ablated. **(D)** Intact bursa of a female, full of sperm, after mating with a male with both aedeagal lateral process tips ablated. Scale bars = 200 µm.

The aedeagus (intromittent organ, phallus, penis) was discovered when anaesthetized copulating pairs were gently pulled apart while submerged in saline solution. This organ in *D. bipectinata* is translucent, membranous and pliable, and it appears to have a textured (scaly) surface (Fig. 5A). Sperm were readily identified emanating from the tip of the aedeagus (Fig 5A, B), and could be gently drawn out of the aedeagus using fine stainless steel minuten pin probes. The aedeagus itself arises from between the bases of the aedeagal lateral processes (Fig. 5B), and was not obviously apparent in any of our KOH-boiled preparations. In our SEMs, a reticulated mat of tissue between the bases of the claws could be discerned (Fig. 3F), which we interpret to be the collapsed aedeagus. In the female of the separated pair, sperm was observed emanating from the female gonopore (the orifice of her reproductive tract through which eggs also exit) (Fig. 5C), and likewise could be pulled further out of the gonopore with minuten pin probes (arrowed Fig. 5D). These observations of sperm emanating simultaneously from the male aedeagus and female gonopore in real time as pairs were gently pulled apart during mating establishes the route of sperm transfer in *D. bipectinata*.

**Figure 5.**
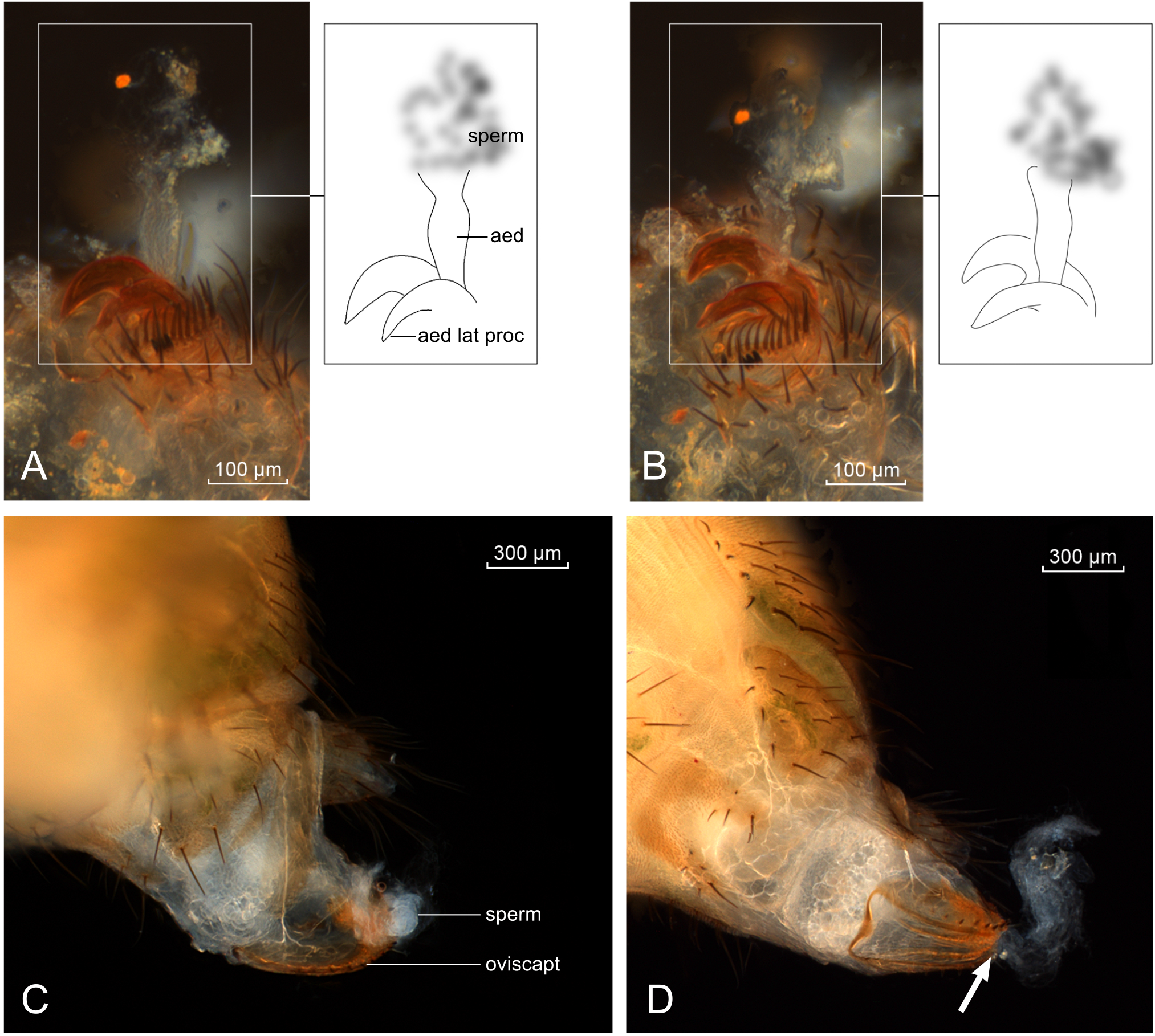
Light microscope images of the genitalia of a male-female pair *D. bipectinata* gently pulled apart during copulation and imaged, demonstrating the extruded aedeagus and female oviscape. **(A), (B)** The aedeagus of the male is clearly shown with sperm emanating from its tip; **(B)** the aedeagus arises from between the bases of the aedeagal lateral processes. **(C)** Sperm simultaneously seeping from the female gonopore, and **(D)** sperm teased and pulled further out from the gonopore (arrowed).

Of the 30 total copulations with “cut” males (those whose claw tips were surgically ablated (Fig. 6)), 22 (73%) resulted in sperm transfer to the bursa (Table 1). In all of these 22 cases, the sperm dissected from the bursa immediately at the end of copulation occurred as a single mass (Fig. 4C, D). Among the males that transferred ejaculate, mean (SE) copulation duration did not differ significantly between cut (10.59 (0.730) min, n = 22) and uncut control (10.12 (1.713) ± 1.77 min, n = 4) (t = 0.25, df = 24, *P* = 0.80) males. Variance in copulation duration for cut males (12.941) was greater than that for uncut males (3.314), but not significantly so (*P* > 0.10). Overall, mean copulation duration between *D. bipectinata* and *D. parabipectinata* did not differ significantly (t = 0.47, df = 33, *P* = 0.64).

**Figure 6.**
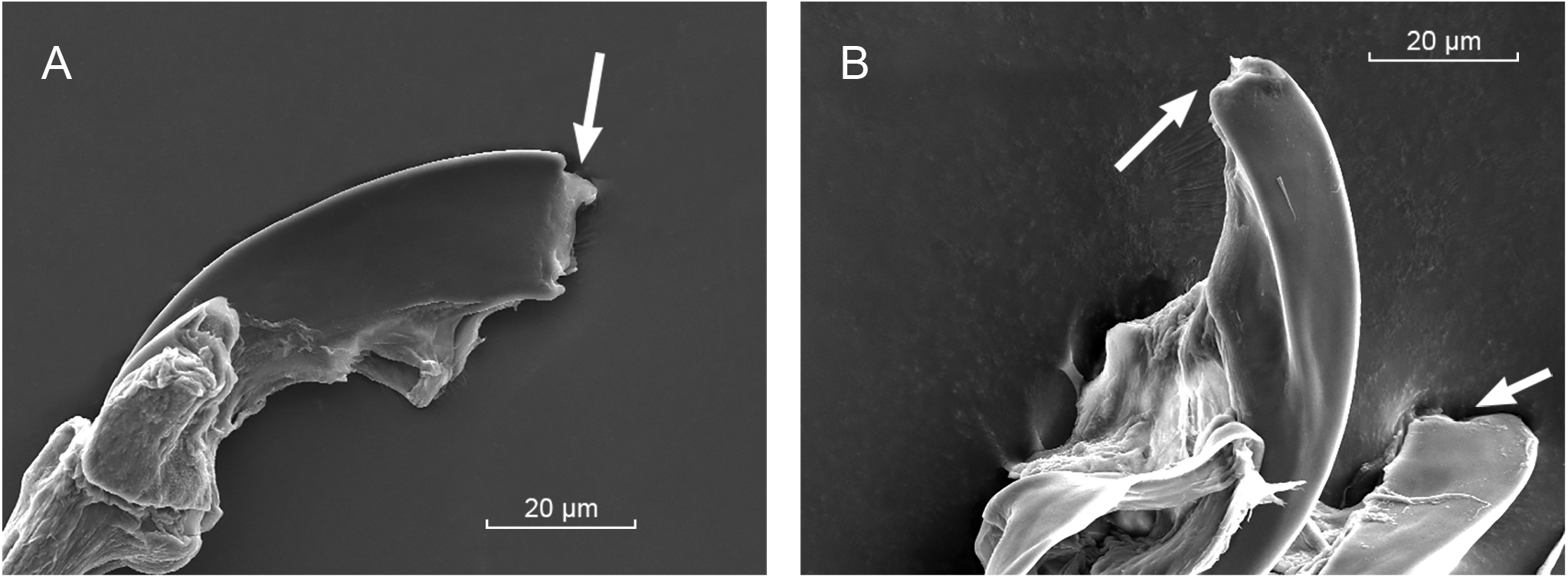
Scanning electron micrographs (1200×) of cut claws, or aedeagal lateral processes in *D. bipectinata*. Examples **(A)** and **(B)** of the processes experimentally blunted (arrowed) using ultra-precise laser surgery are shown. Both tip-ablated processes are visible in **(B)**. From one third to one half of the distal ends of both aedeagal lateral processes of each male were ablated, completely eliminating their sharp terminal ends.

**Table 1.** Number of ejaculates transferred to the female bursa as a single mass in male *D. parabipectinata* and *D. bipectinata*. The aedeagal lateral processes (claws) were cut with an ultraprecise surgical laser; the uncut treatment group consisted of surgical controls. Cut males had approximately 1/3–1/2 the distal ends of both aedeagal lateral processes ablated (Fig. 6). All females were dissected immediately after pairs separated. In all cases of ejaculate transfer, the sperm occurred as a single mass within the female bursa irrespective of whether the male was cut or uncut.

| Species | Surgical treatment | Number pairs set up | Number copulations | Number copulations resulting in ejaculate transfer | Number of ejaculates as a single mass |
| --- | --- | --- | --- | --- | --- |
| *D. parabipectinata* | Cut | 8 | 6 | 5 | 5 |
|  | Uncut | 2 | 2 | 2 | 2 |
| *D. bipectinata* | Cut | 34 | 24 | 17 | 17 |
|  | Uncut | 3 | 3 | 3 | 3 |
| Total | Cut | 42 | 30 | 22 | 22 |
|  | Uncut | 5 | 5 | 5 | 5 |

Among the 8 cut males (out of 30) that copulated but failed to transfer sperm to the female bursa, 2 males were observed to dismount but could not disengage their genitalia from that of the female, and remained fastened to the female in an end-to-end position. What appeared to be ejaculate seeped out from between one pair, and remained attached to the male’s terminalia after the pair finally separated (Fig. 7). This viscous, whitish mass contained sperm, verifying that it was leaked ejaculate.

**Figure 7.**
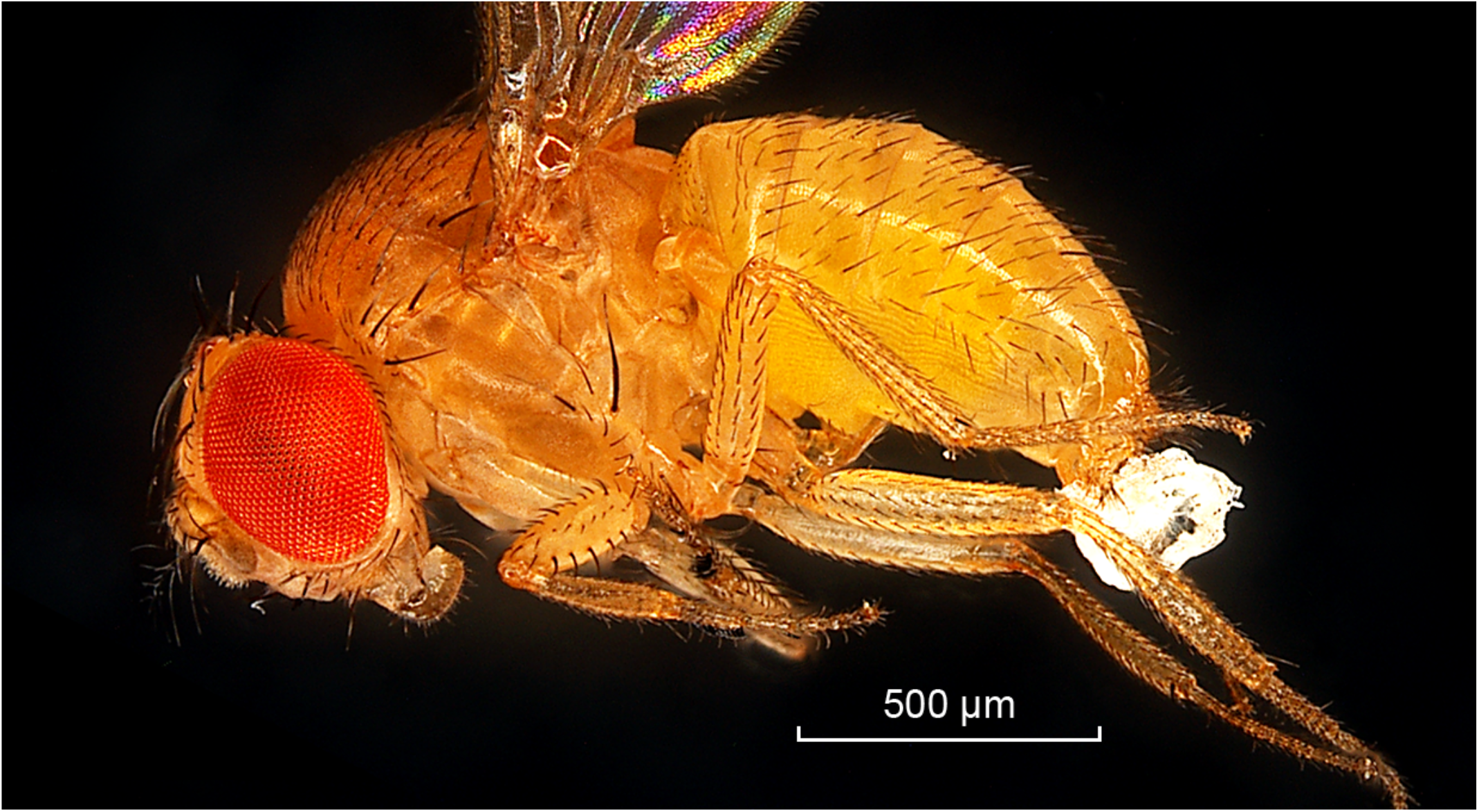
The ejaculate mass, adhered to the tip of a cut male’s abdomen after copulation, had seeped out from between the pair during copulation. The tips of the aedeagal lateral processes in this *D. bipectinata* male had been ablated (Fig. 6) with a surgical laser prior to mating.

## 4. Discussion

In their authoritative review of copulatory wounding, Lange et al. (2013) listed a set of criteria for establishing the existence of traumatic mating in a given species, and here, building upon this work and that of Tatarnic et al. (2014), we assemble a set of criteria for establishing the occurrence of TI. We suggest that evidence for TI should minimally include: *i*) a specific wounding structure(s) that demonstrably breaches the female body wall; *ii*) physical features of said structure(s), such as a canal, lumen, groove and/or pore, for the transfer and delivery of spermatozoa; and *iii*) the transfer of spermatozoa across the female body wall.

Several studies have demonstrated TI by fulfilling these criteria (e.g., Davis, 1956; Carayon, 1966; Řezáč, 2009; Tatarnic and Cassis, 2010), a paradigmatic example of which occurs in bed bugs (Cimicidae) (Carayon, 1966; Usinger, 1966; Benoit, 2011). Copulation in several cimicid species has been observed directly, and it has been unambiguously documented to involve males breaching the female’s body wall with their needle-like “parameres”, often stabbing the female multiple times during a single mating event, and demonstrably transferring sperm into her body cavity (Carayon, 1966). The needle-like paramere, with its readily discernible channel and subterminal pore (Fig. 1), are phenotypic features that reflect the paramere’s function in TI (Siva-Jothy, 2006).

In contrast, we contend that none of the above criteria for demonstrating TI were convincingly met by K2007. In the first place, whereas K2007 claims that integumental penetration is achieved by the claws, stating that they “…pierce the pockets during copulation…” (p. 403), there was no direct evidence presented for physical penetration of the female body wall (nor the genital tract for that matter). The presence of melanized patches in mated females was presented, but this is not decisive evidence for penetration of the integument. Such scarring cannot exclude other possible causes such as surface injury without perforation, and in any event such lesions are known to occur during mating without insemination in other species (Merrit, 1989; Blankenhorn et al., 2002; Kamimura, 2010; Lange et al., 2013). The second criterion (the functional morphology of the organ) was not fulfilled either, as convincing visual evidence for a structure that could guide and transfer sperm across the female body wall was also not provided, and according to the present investigation, does not exist (and see below). Finally, although K2007 claimed that “…sperm is ejaculated through the wounds but not through the genital orifice” (p. 404), pink areas of coloration in a laser scan micrograph were presented as evidence for this claim, which, to us, is insufficient since the presence of sperm within these pink “clouds” was not confirmed, let alone evidence of sperm transfer via the claws to the reproductive tract.

Here, we examined SEMs of the dorsal, lateral and ventral surfaces of the claws in *D. parabipectinata* and *D. bipectinata*, and discerned no obvious “groove” that could transport sperm. We also evaluated the assertion that the paired claws serve to inject sperm into the reproductive tract, by testing the prediction that immediately after and/or during mating, the ejaculatory material within the female should be discernable as two distinct masses. This prediction failed, as sperm invariably occurred as a single mass. This outcome aligns with previous work on the reproductive biology of *D. bipectinata*: in all cases, the sperm dissected from the bursa immediately after copulation occurred as a single mass (Tyler et al., 2021).

The aedeagus (intromittent organ, phallus, penis) in the *bipectinata* complex has been notoriously difficult to detect and characterize. Throughout the genus *Drosophila*, the aedeagus is usually a tubular organ with an external opening, a phallotrema or gonopore. The organ itself is usually membranous, often expanded apically and hirsute or irregularly papillate (Bock and Wheeler, 1972). K2007 suggested that in the *bipectinata* species complex, the aedeagus had diminished to “a degenerate, transparent, tube-like true aedeagus” (p. 403), while Rice et al. (2021) referred to the aedeagus as “translucent” and outlined the area where it ought to be located in *D. malerkotliana*, between the paired lateral processes (figure 2G in Rice et al., 2021). Here, for the first time we have imaged the everted aedeagus (in *D. bipectinata*) through adopting a method of gently separating copulating pairs while under ether anesthesia and submerged in saline solution. Pulling pairs apart during coupling extruded the male aedeagus and clearly showed sperm emanating from its tip and simultaneously from the female vaginal opening (her gonopore). Sperm emanating from these male and female structures in real time identifies the path of sperm transfer to the female, and explains why the sperm mass invariably consists of a single integrated unit, and why males with cut claw tips were able to inseminate females. The aedeagus is a translucent, membranous, and highly pliable tube-like structure that appears to readily collapse upon itself. These characteristics suggest why the aedeagus has been difficult to detect in previous works.

We also tested the prediction of the TI hypothesis that after experimentally eliminating the sharp terminal ends of the claws, insemination should be inhibited. This prediction also failed, as a plurality of males without these sharp ends successfully inseminated females. In matings with ablated males (claw tips removed), 73% resulted in insemination, and in *all* these cases, the sperm occurred as the typical single mass within the female reproductive tract. In all of our experimental males with cut claws, the efficacy of the surgery was verified afterwards; all surgeries had successfully removed the apical third to apical half of both aedeagal lateral processes (claws).

Taken together, the results refute the hypothesis of TI in the *bipectinata* species complex, and therefore in *Drosophila*, and for that matter in Diptera as far as we know. Sperm transfer in the *bipectinata* complex occurs from the male aedeagus into the female reproductive tract via her gonopore, and comports with knowledge about other *Drosophila* species including *D. melanogaster*. An alternative possibility is that the claws may serve to transfer (or secrete) fluid, and therefore function in “traumatic secretion transfer” (Lange et al., 2013), evidence for which occurs in the seed beetle *Callosobruchus maculatus* (Hotzy et al., 2012) and blowfly *Lucilia cuprina* (Merritt, 1989), possibly accounting for the pink areas highlighted in K2007’s images. This idea, however, is speculative, and we have no evidence for or against it.

From our observations of matings in *D. bipectinata* and *D. parabipectinata*, the most likely functions of the claws that we can discern are at least three-fold, all of which are mechanical in nature. We emphasize, however, that additional experiments are needed to fully characterize the function of these remarkable structures, but which are beyond the scope of the present study. Here, our primary focus was to address the TI hypothesis in and of itself.

A first potential function we may deduce from our data is that the claws serve to assist the male in achieving copulation by facilitating the grasping of the female. In 10 out of 30 cases of copulations with cut males, males were observed to mount the female, probe the female terminalia with their own genitalia, but failed to achieve genital coupling in these attempts (some later did). A grasping function has been demonstrated for the sharp (periphallic) spines emanating from the male ventral cercal lobes in *D. bipectinata* (Polak and Rashed, 2010) and *D. ananassae* (Grieshop and Polak, 2012). More generally, male genital clasping devices, such as spines, hooks, inflatable organs/structures, and other interlocking features, occur in a wide range of invertebrate species to function in achieving union and genital integration, holding the female securely, and protecting her from rival males (Thornhill and Alcock, 1983; Eberhard, 1985; Gwynne, 1998; Simmons, 2001).

A second apparent function may be to assist in the opening of the female gonopore. In 2 of the above 10 instances of cut males failing to couple, males were observed to use their periphallic structures (surstyli or claspers) to probe and apparently attempt to part the female ovipositor, but failed to do so, and failed to achieve union. These observations suggest that the female ovipositor could not be opened as a result of the males lacking intact claws.

A third non-mutually exclusive function we believe is one for anchoring—to brace the male genitalia to allow the intromittent organ (aedeagus) to evert through the female gonopore into the vagina. A firm integration of the genitalia should assure transfer of ejaculate from the aedeagus to the female. Passage of the aedeagus through the female gonopore could also be facilitated or provided by the physical opening of the gonopore by the abduction of the claws. A relevant observation here is that there were 2 cases (out of 8) in which cut males remained “adhered” to the female after dismounting, suggesting that ejaculate failed to transfer to the female because of poor union of the genitalia, seeped from between the pair, and acted as an adhesive that then unnaturally prolonged genital coupling. Sperm seepage is consistent with clawless males being unable to adequately insert the aedeagus into the gonopore and/or effectively maintain their genitalia integrated with that of the female during copulation. Ejaculate seepage during coupling was directly observed in one case (Fig. 7).

When TI occurs in a given species, it is predicted to impose a number of potential costs to females, some of which will be unique relative to other forms of copulatory wounding. In cases of TI, we may expect females to incur specific fitness costs associated not only with wound healing, immune system activation and risk of infection of the hemocoel and internal organs, but also with the physiological challenges stemming from the introduction of seminal fluid and spermatozoa into the hemolymph (Davis, 1956; Morrow and Arnqvist, 2003; Siva-Jothy, 2009). This latter cost could also encompass loss of control over fertilization that would normally be available to females where insemination occurs via the female reproductive tract (Eberhard, 1996; Beani et al., 2005). Our ability to predict such costs to females, and to interpret aspects of female reproductive anatomy, physiology, and behaviour, that evolve in response to sexual conflict linked to mating (Lessels, 2006; Parker, 2006; Yassin and Orgogozo, 2013), rests upon accurately describing and classifying the highly varied forms of copulatory trauma that exist in animals. The results of the present study lead us to reject the assertion that TI occurs in species of the *Drosophila bipectinata* complex.

## Acknowledgements

We thank Melodie Fickenscher of the Advanced Materials Characterization Center at University of Cincinnati for her expertise in taking the SEMs, and Joshua B. Benoit for the bed bug specimen and for helpful discussions.

